# Association between Apolipoprotein E genotype and the gut microbiome composition in humans and mice

**DOI:** 10.1101/473694

**Authors:** Tam T.T. Tran, Simone Corsini, Lee Kellingray, Claire Hegarty, Gwénaëlle Le Gall, Arjan Narbad, Michael Müller, Noemi Tejera, Paul W. O’Toole, Anne-Marie Minihane, David Vauzour

## Abstract

Apolipoprotein E (*APOE*) genotype is the strongest prevalent genetic risk factor for Alzheimer’s disease. Numerous studies have provided insights into the pathological mechanisms. However, a comprehensive understanding of the impact of *APOE* genotype on microflora speciation and metabolism is completely lacking. The aim of this study was to investigate the association between *APOE* genotype and the gut microbiome composition in human and *APOE-*targeted replacement (TR, *APOE3* and *APOE4*) transgenic mice. Faecal microbiota amplicon sequencing from matched individuals with different *APOE* genotypes revealed no significant differences in overall microbiota diversity (alpha or beta diversity) in group-aggregated human *APOE* genotypes. However, several bacterial taxa showed significantly different relative abundance between *APOE* genotypes. Notably, we detected an association of *Prevotellaceae* and *Ruminococcaceae* and several butyrate-producing genera abundances with *APOE* genotypes. These findings were confirmed by comparing the gut microbiota of *APOE*-TR mice. Furthermore, metabolomic analysis of faecal water from murine samples detected significant differences in microbe-associated amino acids and short-chain fatty acids between *APOE* genotypes. Together, the findings indicate that *APOE* genotype associated with specific gut microbiome profiles in both humans and in *APOE*-TR mice. This suggests that the gut microbiome is worth further investigation as a potential therapeutic target to mitigate the deleterious impact of the *APOE4* allele on cognitive decline and the prevention and treatment of AD.

## Introduction

The gut microbiome is intimately involved in numerous aspects of human physiology. Emerging evidence links perturbations in the microbiome to neurodegeneration and Alzheimer’s disease (AD), with (neuro)inflammation proposed as an aetiological link (Tremlett *et al*., 2017).

The extent to which host genetic variation determines the microbiome composition is still currently debated. Indeed, although previous studies have reported that the microbiomes of humans and mice are associated with host genetic variation (Spor *et al*., 2011) and have identified several heritable bacterial taxa (Beaumont *et al*., 2016; Bonder *et al*., 2016; Turpin *et al*., 2016), other studies have identified a stronger environmental influence compared to host genetics in shaping human gut microbiota (Rothschild *et al*., 2018). Thus, the extent to which human genetics shape microbiome composition remains unclear.

*Apolipoprotein* E (*APOE*) genotype is the strongest prevalent risk factor for neuropathology and AD (Liu *et al*., 2013; Neu *et al*., 2017; Pontifex *et al*., 2018). ApoE was originally identified as a component of systemic circulating lipoproteins and a member of a family of apolipoprotein modulators of their metabolism. It has subsequently emerged as the almost exclusive lipid transporter in the central nervous system (Shore and Shore, 1973; Vauzour and Minihane, 2012). In humans, APOE exists in three different isoforms (apoE2, apoE3 and apoE4), arising from three different alleles (ε2, ε3 and ε4). These alleles give rise to three homozygous (*APOE2/E2, APOE3/E3* and *APOE4/E4*) and three heterozygous (*APOE3/E2*, *APOE4/E3* and *APOE4/E2*) genotypes in humans (Huang *et al*., 2004). Generally, 50-70% of populations present with the *APOE3/E3* genotype, with the ε3 allele accounting for 70-80% of the gene pool, and the ε2 and ε4 allele accounting for 5-10% and 10-15% respectively (Huang *et al*., 2004). *APOE4* carrier status is highly predictive of dementia and AD, with *APOE3/E4* and *APOE4/E4* being at 3-4 and 8-12 fold increased risk and a much earlier age of onset (Pontifex *et al*., 2018). Although the aetiological basis of *APOE4-*neuropathological associations has been widely researched and reported, the main aetiological mechanism has not been clearly defined. The ApoE protein is involved in multiple biological processes including lipoprotein metabolism (Raffai *et al*., 2001), intracellular cholesterol utilisation (Reyland and Williams, 1991), cell growth (Ishigami *et al*., 1998), immunoregulation, (neuro)inflammation (Hui *et al*., 1980; Pepe and Curtiss, 1986) and neuroprotection (Jofre-Monseny *et al*., 2008). Whilst involved in the metabolism of gut-derived chylomicrons, its role in intestinal integrity and gut microbiome composition and metabolism is currently unknown.

In the present study, we explore the hypothesis that *APOE* variation influences the microbiome composition and its subsequent metabolism. Our experiments using human faecal samples and *APOE*‐ targeted replacement (TR) mice revealed significantly different relative abundance between bacterial taxa according to *APOE* genotypes. Furthermore, using a metabolomic approach, differences in microbe-associated amino acids and short-chain fatty acids according to *APOE* genotypes were also observed. Taken together, our findings indicate that *APOE* genotype associates with specific gut microbiome profiles, which may affect the host metabolism and ultimately contribute to AD pathology.

## Results

### Descriptive statistics of human *APOE*-genotyped cohorts

A total of 56 fecal samples were analysed from participants of the COB (NCT01922869) and the CANN studies (NCT02525198) (Norwich Clinical Centre, UK) with the four *APOE* genotype groups selected to be matched for sex, age and body mass index (BMI) (**Table 1** and **Table S1**).

**Table 1.** Clinical characteristics of participants according to *APOE* genotypes.

| Parameter | *E2/E3* | *E3/E3* | *E3/E4* | *E4/E4* | *p*-value<sup>†</sup> |
| --- | --- | --- | --- | --- | --- |
| n | 14 | 18 | 18 | 6 |  |
| Age (yrs) | 68.6 ± 4.6 | 68.5 ± 5.0 | 68.6 ± 3.0 | 67.7 ± 6.1 | 1.0 |
| Sex (# male:female) | 7:7 | 9:9 | 9:9 | 3:3 | 1.0 |
| BMI (kg/m<sup>2</sup>) | 25 ± 2.2 | 26.3 ± 2.6 | 26.1 ± 3.1 | 25 ± 1.9 | 0.42 |

Data presented as mean ± SD. <sup>†</sup>*p*-value was calculated by Kruskal-Wallis *H* test.

### Difference in human gut microbiota composition between *APOE* genotypes

The V3-V4 hypervariable region of the 16S rRNA gene was PCR-amplified from faecal samples collected from participants to generate an amplicon of approximately 460 bp. Sequencing this amplicon allows determination of microbiota composition. The reads were clustered at a 97% similarity threshold into 3,314 unique Operational Taxonomic Units (OTUs) or sequence-based bacterial classification, approximating to species. The total OTUs were assigned to 15 phyla, 27 classes, 43 orders, 70 families, and 155 unique genera, across the entire dataset. The vast majority (99.5 %) of all sequences were affiliated to five dominant phyla, mainly in the *Firmicutes* (82.2±10.8%), lower assignment to phyla *Bacteroidetes* (7.7±6.3%), *Actinobacteria* (3.8±4.5%), *Proteobacteria* (3.2±9.5%), and *Verrucomicrobia* (2.6±5.1%) (**Figure S1a)**. After rarefaction with a depth of 8,736 reads per sample, alpha diversity (net diversity within a single sample/subject) was measured by calculating three diversity indices namely chao1 (richness), phylogenetic diversity, and Shannon diversity index. None of these metrics was significantly different between *APOE* genotypes (**Figure S2a**). Similarly, there was no significant difference in any of the alpha diversity-metrics between males and females (**Figure S2b**). However, the microbiota alpha diversity of obese subjects (n=3) was significantly lower than that in normal weight and overweight subjects (p < 0.05, **Figure S2c**), in line with previous observations (Le Chatelier *et al*., 2013). Beta-diversity analysis (which measures inter-individual microbiota relatedness) was performed using Principal Coordinate Analysis (PCoA) clustering based on unweighted and weighted UniFrac distances. PERMANOVA for testing associations between clinical parameters and microbiota composition are given in **Table S2**. There was no difference in beta diversity of gut microbiota composition according to *APOE* genotype (**Figure S3a**). However, we observed a weak, but significant association between microbiota composition and gender and BMI categories (**Figure S3b** and **Figure S3c**).

Although alpha and beta-diversity analyses of the gut microbiota did not discriminate between *APOE* genotypes, these are global measures that detect relatively large differences in microbiota structure. We therefore questioned whether the relative abundance of any taxa might differ between these genotypes, using Kruskal-Wallis H test to compare all taxa at various phylogenetic assignment levels across all genotypes. We observed that the relative abundance of the phylum *Firmicutes* and order *Clostridiales* was higher in subjects of the *APOE2/E3* genotype than in *APOE3/E4* or *APOE4/E4* (*p* < 0.05; **Figure 1** and **Table S3**). Furthermore, at the bacterial family level, the abundance of *Ruminococcaceae* (a family of fermentative anaerobes associated with fibre degradation and short chain fatty acid production) was higher in *APOE2/E3* than in *APOE3/E3* (*p* = 0.004), *APOE3/E4* (*p* = 0.002) or *APOE4/E4* (*p* = 0.072). On the other hand, the abundance of *Prevotellaceae* was lower in *APOE2/E3* than the other three *APOE* genotypes (*APOE3/E3 p* = 0.008, *APOE3/E4 p* = 0.085, *APOE4/E4 p* = 0.015) and was slightly more abundant at close to significant levels (*p* = 0.088) in *APOE3/E4* compared to *APOE4/E4* with mean of relative abundance of 1.79% versus 1.40% (**Figure 1c**, **Table S3** and **Figure S1b)**. Within the *Ruminococcaceae* family, three genera including *Clostridium* cluster IV*, Clostridium* cluster XIVa and *Gemmiger* were statistically significant and differentially abundant according to *APOE* genotypes. The abundance of *Clostridium* cluster IV was lower in *APOE3/E3* than in *APOE2/E3* (*p* = 0.027) and *APOE4/E4* (*p* = 0.039) while the abundance of *Clostridium* cluster XIVa was higher in *APOE4/E4* than in *APOE2/E3* (*p* = 0.044) and *APOE3/E4* (*p* = 0.078). Higher presence of *Gemmiger* was observed in fecal samples from *APOE2/E3* compared to *APOE3/E3* (*p* = 0.0499) and *APOE3/E4* (*p* = 0.086). Moreover, we observed a higher abundance of *Roseburia* in fecal samples at close to significant levels (*p*< 0.1) in *APOE3/E3* compared to three other *APOE* genotypes and in *APOE3/E4* compared to *APOE4/E4* (**Figure 1d, Table S3** and **Figure S1c).**

**Figure 1.**
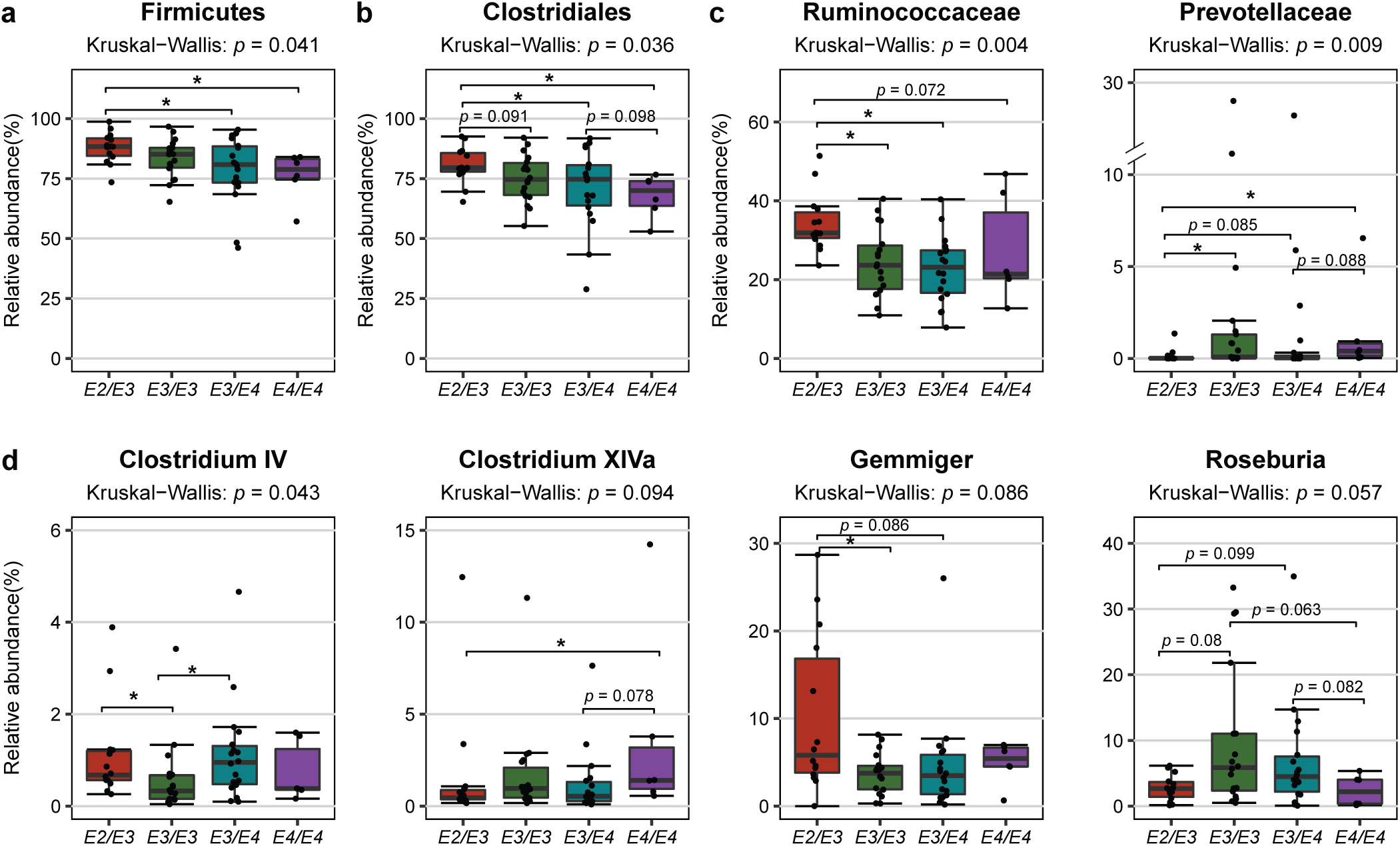
Box plot of the relative abundance distribution of selected taxa at (**a**) Phylum level, (**b**) Order level, (**c**) Family level and (**d**) Clostridium cluster/Genus level. Taxa were selected based on differential abundance associated with human *APOE* genotypes. *P*-values were calculated by Kruskal-Wallis H test for all genotypes with followed by Dunn’s multiple comparisons adjusting false discovery rate using the Benjamini-Hochberg correction, \**p* < 0.05.

To determine possible associations between *APOE* genotypes and microbial translocation, we measured the plasma levels of two biomarkers of intestinal integrity, namely haptoglobin (Hp) and lipopolysaccharide binding protein (LBP). No significant differences were observed in the levels of Hp and LBP (**Table S1**) according to genotype. Furthermore, no significant correlation was observed between the clinical parameters and both the weighted and unweighted Unifrac distances (**Table S2**).

### Difference in murine gut microbiota composition between *APOE* genotypes

We next sought to investigate if *APOE* genotype/gut microbiota interactions in humans were evident in human transgenic homozygous *APOE3-* and *APOE4-*TR mice at 4 months (young) and 18 months (old) of age. Their gut microbial communities were analyzed based on sequencing the V4 hypervariable region (approximately 254 bp) of the 16S rRNA gene. There was no significant difference in alpha diversity between *APOE3* and *APOE4* genotypes. However, in line with our previous studies of microbiota in ageing humans (O’Toole and Jeffery, 2015) and rodents (Flemer *et al*., 2017), both chao1 and phylogenetic diversity were much higher in young mice compared to old mice (*p* < 0.001; **Figure S4**). Moreover, UniFrac distances (unweighted and weighted) PCoA showed that faecal microbial profiles of young mice separated significantly from those of old mice (PERMANOVA, *p* = 0.001; **Figure 2a**). Within each age group, both UniFrac measures showed significant microbiota differences between *APOE3* and *APOE4* genotypes, with the *p*-value from PERMANOVA analysis less than 0.005 (**Figure 2a**). These differences could be explained by differences detected in the relative abundance of dominant taxa, of which the most dominant were *Firmicutes* (62.8 ± 14.4%) and *Bacteroidetes* (32.3 ± 14.8%), followed by *Proteobacteria*, *Verrucomicrobia* and *Deferribacteres* accounting for less than 5% in total (**Figure S5a)**.

**Figure 2.**
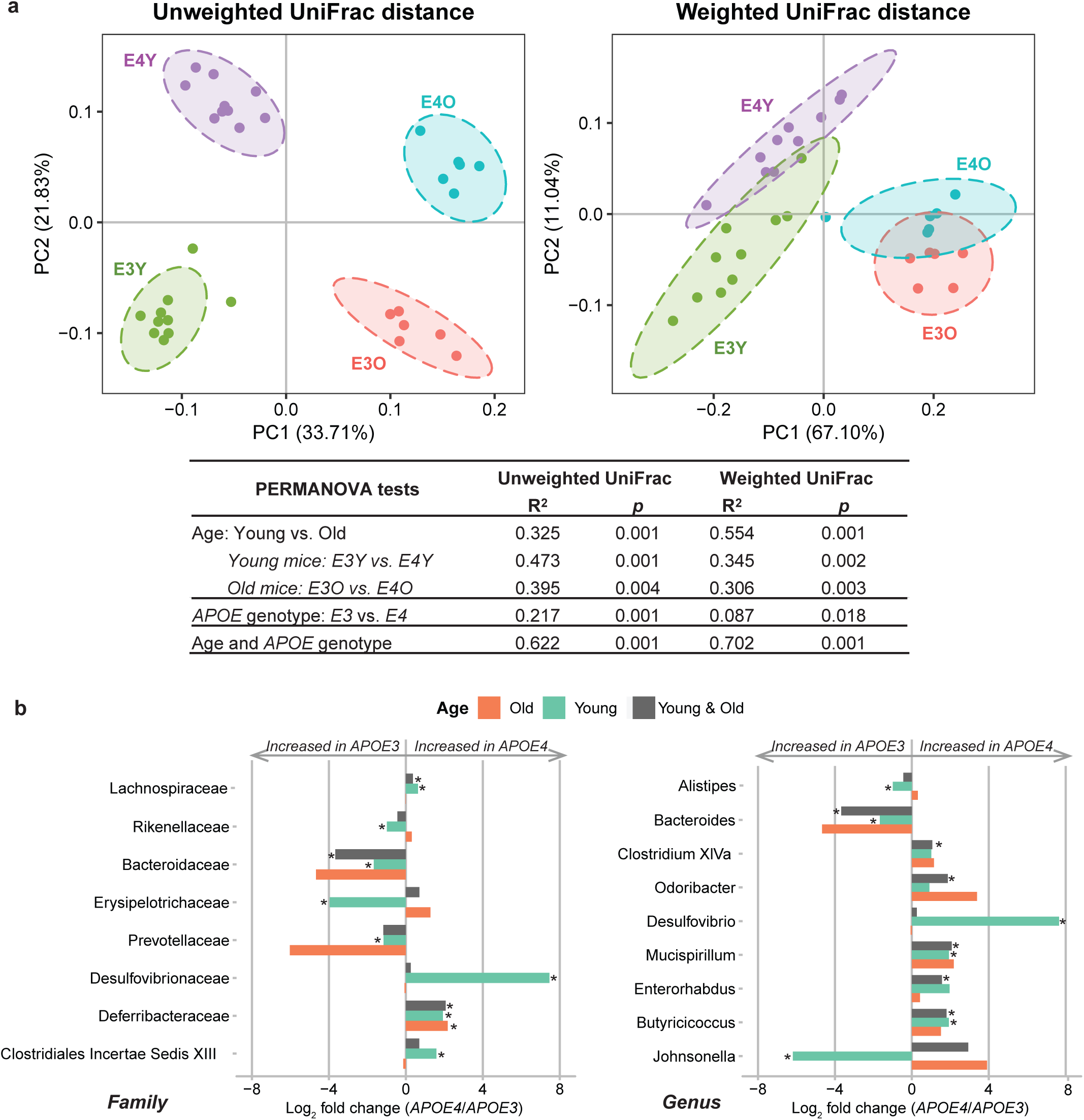
Differences in gut microbiome composition between *APOE3*-TR and *APOE4*-TR mice. (**a**) Principle Coordinates Analysis based on unweighted and weighted UniFrac distances of partial sequences of bacterial 16S rRNA genes showing gut microbiota beta diversity grouped by age and *APOE* genotypes. Significant differences between groups were calculated by PERMANOVA tests. (**b**) Comparison of relative abundance taxa between *APOE3* and *APOE4* in young mice samples, old mice samples and both age groups combined were represented by log_2_fold changes. Statistical significances were determined by the Mann–Whitney U test and were corrected for the multiple comparison using the Benjamini–Hochberg adjustment, \**p* < 0.05. E3Y, *APOE3* young mice; E4Y, *APOE4* young mice; E3O, *APOE3* old mice; E4O, *APOE4* old mice.

Analysis of differentially abundant taxa between *APOE3*-TR and *APOE4*-TR animals at the phylum level revealed that *Deferribacteres* in combined young and old mice were notably higher in the *APOE4*-TR mice compared to the *APOE3*-TR mice, while the opposite was true for *Candidatus Saccharibacteria*. In addition, lower relative abundance of *Proteobacteria* were seen in *APOE4*-TR young mice when compared to the *APOE3*-TR young mice **(Table S4** and **Figure S6)**. Although no significant difference was found in aggregated *Firmicute* or *Bacteroidetes* phylum abundance between *APOE* genotypes, we observed an increase in *Firmicutes*/*Bacteroidetes* ratio in old mice when compared to young mice (p < 0.001, **Figure S7**), in agreement with a previous C57BL/6N mouse study (Hoffman *et al*., 2017). At the order level, *Deferribacterales* abundance in combined age groups was significantly higher in the *APOE4*-TR mice compared to the *APOE3*-TR mice. Additionally, *Clostridiales*, *Erysipelotrichales* and *Desulfovibrionales* in young mice were significantly different in relative abundance between the two *APOE* genotypes. The increase of *Lachnospiraceae*, *Deferribacteraceae* abundance and decrease of *Bacteroidaceae* abundance at family level in *APOE4* transgenes compared with *APOE3* was detected in combined age groups. *Desulfovibrionaceae*, *Clostridiales Incertae Sedis* XIII, *Rikenellaceae*, *Prevotellaceae*, *Erysipelotrichaceae* were also found to be significantly different between *APOE* genotypes in young mice **(Figure 2b, Table S4**, and **Figure S6)**. Those differentially abundant families by *APOE* genotype were reflected by *Mucispirillum*, *Clostridium* cluster XIVa, *Butyricicoccus*, *Odorobacter*, *Enterorhabdus*, and *Bacteroides* in combined age groups; *Mucispirillum*, *Desulfovibrio*, *Butyricicoccus*, *Bacteroides*, *Alistipes* and *Johnsonella* in young mice at genus level **(Figure 2b, Table S4**, and **Figure S6)**.

### Faecal metabolite associations with *APOE* genotype and age

In order to improve our understanding of the relationships between metabolite and microbiota composition in the gut, we performed metabolomic analyses of faecal water prepared from caecal contents. Sparse PLS Discriminant Analysis (sPLS-DA) showed a trend for separation according to age and *APOE* genotypes **(Fig S8)**. Two-way ANOVA was therefore performed to investigate interactions between age and *APOE* genotype. Seven metabolites, adenosine monophosphate, alpha ketoisovaleric acid, glucose, glycine, lactate, oxocaproate, and xanthine were present at significantly different levels in age-*APOE* genotype interaction while 39 and 19 metabolites were significantly different in age groups and *APOE* genotype groups, respectively (**Figure S9, Table S5**). Four clusters of all significant metabolites had distinct correlations. Cluster A comprising 5 metabolites (lactate, pyruvate, fumarate, hypoxanthine and uracil) had inverse direct correlations with *APOE4*-TR old mice and had strong direct correlations with three other groups. However, cluster B and cluster C metabolites were associated with age. Ten metabolites in cluster B (methylamine, acetate, butyrate, propionate, arabinose, xylose, succinate, glucose, AMP, GTP) were more abundant in young mice especially in *APOE3*-TR young compared to old mice. Fourteen metabolites in Cluster C (asparagine, alanine, tryptophan, threonine, tyrosine, lysine, phenylalanine, glutamate, histidine, leucine, glutamine, valine, isoleucine, methionine) showed an opposite trend. Cluster D metabolites were divided into two sub-clusters, cluster D1 comprising 4 metabolites (2-oxoisocaproate, alpha-ketoisovalerate, 3-methyl-2-oxovalerate, urocanate) had direct correlations with *APOE4*-TR young mice; and cluster D2 including 14 metabolites (isobutyrate, 1,3-dihydroxyacetone, lactaldehyde, aspartate, ornithine, ribose, xanthine, choline, glycine, creatine, taurine, 2-methylbutyric acid, ethanol and formate) which had positive correlations with old mice (**Figure 3a**). In addition, Metabolite Set Enrichment Analysis (MESA) was used to identify significantly enriched pathways in metabolomics data associated with *APOE* genotype and age. Of the top 50 assigned pathways, the significant pathways in *APOE* genotype were ammonia recycling, urea cycle, and alanine metabolism (**Table S6, Figure S10a**) while the significant pathways in age were ammonia recycling, urea cycle, glycine and serine metabolism, glutamate metabolism and alanine metabolism (**Table S7, Figure S10b**).

**Figure 3.**
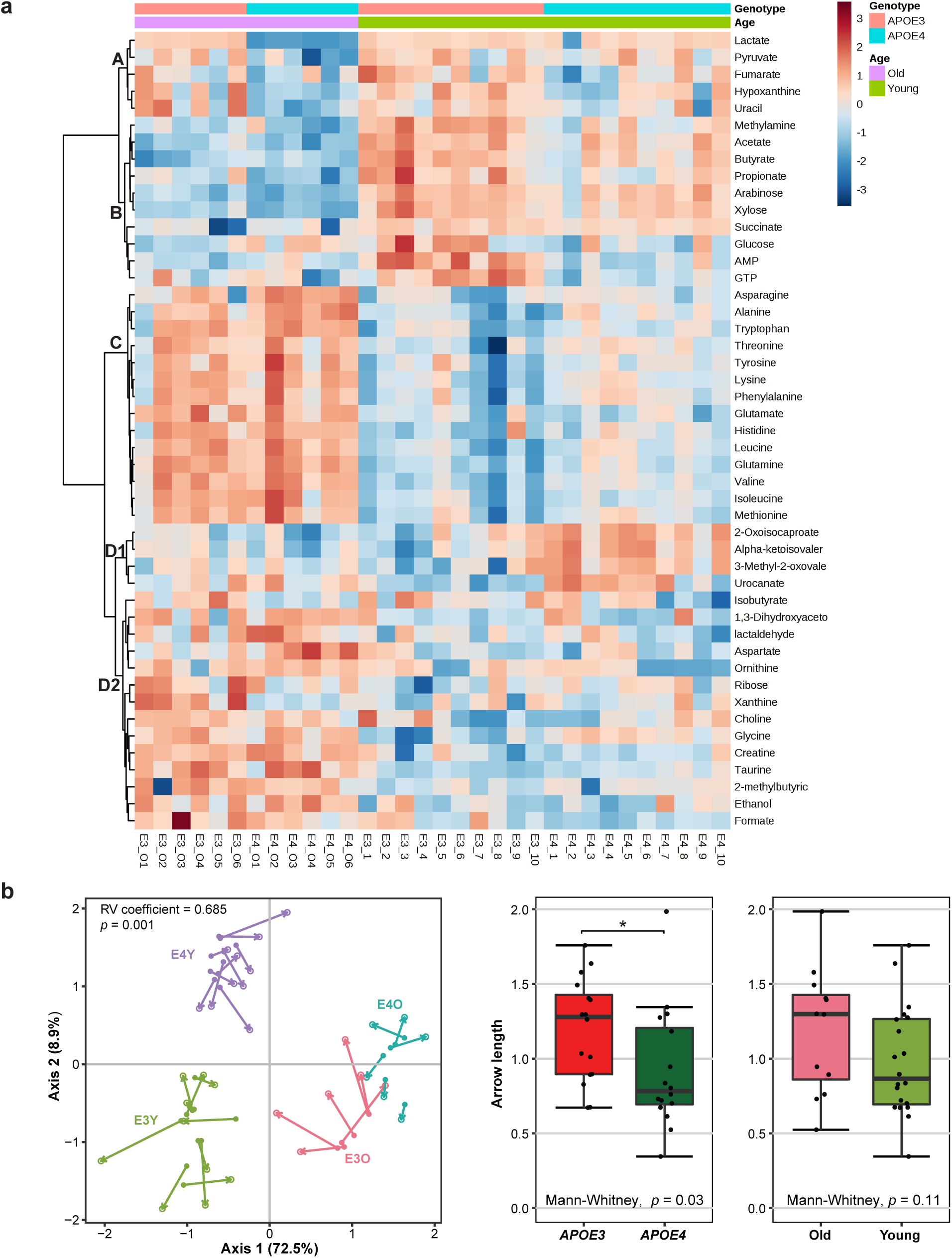
Faecal metabolome analysis of *APOE3*-TR and *APOE4*-TR mice. (**a**) Heatmap and cluster analysis of two-way ANOVA of significantly differentially abundant metabolites grouped by age and *APOE* genotype. Clustering was obtained following similarity analysis using the Ward hierarchical algorithm and Euclidean distance metrics. (**b**) Co-inertia analysis (COIA) of the association between metabolites and microbiota composition in the gut. The left panel shows the COIA of the microbiota principal component analysis (PCA) (solid circle) at OTU level and the metabolomics PCA (empty circle); length of arrow indicates the divergence between two datasets. The right panel shows co-inertia of metabolome and microbiota data, represented by arrow length between the two data points per sample, grouped according to *APOE* genotype or age. Length of arrow was estimated using Euclidean distance measurement. \**p* < 0.05.

Co-inertia analysis (COIA) was carried to explore the correlation between the composition of microbiota at OTU level and the faecal metabolome (**Figure 3b**). The Monte-Carlo permutation test revealed a high overall similarity in the structure between the two datasets which was statistically significant (RV co-efficient = 0.685; *p* = 0.01). The first four axes represented 72.5 %, 8.9%, 7.0% and 2.6% of the explained variance respectively, and so the analysis focused on the first axis. Each sample is represented by an arrow where length of arrow indicates the divergence between two datasets. We observed that the aggregate arrow length was shorter in *APOE4* mice compared with that in *APOE3* mice, which indicated a higher consensus between microbiota composition and metabolites of *APOE4* mice compared with *APOE3* mice. The metabolites and OTUs that strongly correlated in the co-inertia analysis axes were plotted in on the first two COIA axes (**Figure S11)**. Metabolites and bacterial OTUs were projected onto the same direction as samples, indicating that they were more abundant in those samples. There was an agreement between the metabolite abundance and the specific taxon abundance. Notably, SCFAs including acetate, butyrate and propionate were located in the direction of butyrate-producing bacteria from *Clostridium* cluster IV genus and the families *Ruminococcaceae* and *Lachnospiraceae*.

## Discussion

Although several recent studies have implicated a link between the gut microbiome and the development of AD (Bhattacharjee and Lukiw, 2013; Ghaisas *et al*., 2016; Sampson *et al*., 2016; Vogt *et al*., 2017), there is no direct study that establishes a link between gut microbiota composition and the strongest genetic risk factor for AD, *APOE* genotype. The current study marks the first analysis that compares gut microbiota composition in humans and transgenic mice with different *APOE* genotypes. Analysis of 16S rRNA gene sequences and faecal metabolome showed that *APOE* genotype correlated with abundance differences of several gut bacterial taxa which may drive the difference in amino acids and SCFAs levels.

Higher levels of *Prevotellaceae* were evident in *APOE3/E3* carriers relative to other genotype subgroups, while higher levels of *Ruminococcaceae* were correlated with the *APOE2/E3* genotype (**Figure 1 and Table S3**) relative to *APOE4* carriers. Similar associations were observed in *APOE*-TR transgenic animals with an increase in *Prevotellaceae* abundance in *APOE3* young mice compared to *APOE4* young mice in this study and an increase of *Ruminococcaceae* in *APOE2* mice compared to *APOE4* and *APOE3* mice (Parikh *et al*., 2017). Interestingly, loss of these bacteria has been reported to negatively correlate with neurodegenerative disorders. *Prevotellaceae* and *Ruminococcaceae* were noted as being less abundant in patients with Parkinson’s disease (PD) (Unger *et al*., 2016) and AD (Vogt *et al*., 2017). A reduction of *Prevotellaceae* influenced mucin synthesis and increased mucosal permeability, allowing local and systemic exposure to bacterial endotoxin which may lead to the accumulation of alpha-synuclein (α-syn) in the colon (Forsyth *et al*., 2011; Scheperjans *et al*., 2015). Aggregation-prone proteins such as β-amyloid (Aβ) and α-syn can propagate from the gut to the brain via the vagus nerve (Holmqvist *et al*., 2014) and contribute to the pathogenesis of PD, AD, and other neurodegenerative disorders (Bennett, 2005; Crews *et al*., 2009; Jellinger, 2003; Stefanis, 2012). *Ruminococcaceae* are involved in the production of short chain fatty acids (SCFAs), such that their depletion is causally linked to inflammation (Larsen *et al*., 2010; Pryde *et al*., 2002; Zhang *et al*., 2016). These findings suggest that these bacteria might contribute to the protective effects of *APOE2* and *APOE3* alleles against AD relative to the *APOE4* genotype (de-Almada *et al*., 2012; Farrer *et al*., 1997; Liu *et al*., 2013).

The high abundance of *Ruminococcaceae* in subjects of the *APOE2/E3* genotype was reflected by *Gemmiger* at the genus level. *Gemmiger* are strictly anaerobic bacteria which ferment a variety of carbohydrates to produce formic and n-butyric acids, often with small amounts of acetic, lactic, succinic, malonic, and pyruvic acids (Gossling and Moore, 1975). In addition, we observed differences between *APOE* genotypes in *Clostridium* cluster IV, *Roseburia* and *Clostridium* cluster XIVa which are able to convert dietary fibres to SCFAs (Duncan *et al*., 2002; Van den Abbeele *et al*., 2013). Although the butyrate-producing bacteria *Clostridium* cluster IV were modestly less abundant in human *APO*E3/E3 individuals, they could be substituted by *Roseburia* with an increased abundance of these bacteria in the *APOE*3/E3. A slight increase of *Clostridium* cluster XIVa from *Lachnospiraceae* was seen in the human *APOE4/E4* genotype which is consistent with the murine data. However, the increase of this genus in *APOE4/E4* may not substitute for the reduction of other butyrate-producing bacteria. Additionally, several genera, which were not correlated with *APOE* genotype in human gut microbiomes, were significantly different between *APOE3* and *APOE4* genotypes in murine gut microbiomes. These taxa could not be detected in human data due to: (*i*) absence of some mouse gut microbiota in human such as *Mucispirillum*, (*ii*) differences in relative abundance of each individual taxa, and (*iii*) complexity in interactions of human gut microbiota with genetics, diet, and other environmental factors.

Incorporating the gut microbiota data with the corresponding metabolites, faecal samples between *APOE* genotype groups were clearly discriminated based on the metabolomic profiles of faecal water extracts. We detected a higher level of SCFAs including acetate, lactate, propionate, and pyruvate in *APOE3* mice which might be the result of a high abundance of butyrate-producing bacteria. Several SCFAs have been shown to inhibit the formation of toxic soluble Aβ aggregates *in vitro* (Ho *et al*., 2018). Interestingly, co-segregation of the faecal metabolomic profiles and the gut microbiome profiles as revealed by the co-inertia analysis suggests that the differences in gut microbiota associated with *APOE* genotype and age in *APOE*-TR mice are reflected in the segregation of metabolites, which may be clinically relevant.

In conclusion, the collective evidence here suggests a link between *APOE* genotypes and gut microbiome composition. Loss of butyrate-producing bacteria and SCFAs in *APOE4* carriers might drive the impact of the *APOE4* allele on neuropathology. Our findings suggest a possible role of gut microbiota, especially butyrate-producing bacteria, as an intervention point to mitigate the impact of *APOE* genotype in the development of AD.

## Methods

### Participants, sample collection, *APOE* genotyping and biochemical analysis

Fifty-six healthy participants, aged between 56 and 78 years, were prospectively selected according to *APOE* genotype from the COB (NCT01922869) and the CANN (NCT02525198) studies for the analysis of their gut microbiota speciation. Participants were provided with faecal collection kits, which included a stool collection bag and an ice pack. They were asked to defecate directly into the bag, which was secured and placed with the ice pack into an insulated container and delivered to the study scientist. The samples were then homogenised by physical manipulation before aliquots were taken and stored at −80°C. The study protocols were approved by the National Research Ethics Service (NRES) Committee (13/EE/0066 (COB) and 14/EE/0189 (CANN) studies), and all participants consented to provide stool samples, and to the use of the stored samples for research purposes.

*APOE* genotyping was carried out as previously described (Calabuig-Navarro *et al*., 2014). Briefly, DNA was isolated from the buffy coat layer of 8 mL of blood collected into sodium heparin CPT™ Mononuclear Cell Preparation Tubes with the use of the QIAgen DNA blood mini kit (Qiagen Ltd). Allelic discrimination of the *APOE* gene variants was conducted with TaqMan PCR technology (7500 Instrument; Applied Biosystems) and Assay-on-Demand single nucleotide polymorphism genotyping assays (Applied Biosystems). The *APOE* haplotypes (E2/E3, E3/E3, E3/E4, and E4/E4) were determined from the alleles for the *APOE* single nucleotide polymorphisms rs7412 and rs429358. Twenty-four participants were selected as *APOE4* carriers (*APOE*3/4 and *APOE*4/4; 12 men and 12 women), with 32 participants selected as *APOE4* non-carriers (*APOE*2/3 and *APOE*3/3; 16 men and 16 women), with the selection process matching the genotype groups for age, BMI and gender.

Serum LPS Binding Protein (LBP) (cat. Ab213805, Abcam, Cambridge, UK) and Haptoglobin (cat. ab108856, Abcam, Cambridge, UK) plasmatic concentrations were detected by enzyme-linked immunosorbent assay (ELISA) kits according to the manufacturer’s instructions. The assay range for the LBP and the Haptoglobin ELISA kits were 1.56 ng/ml – 100 ng/ml and 0.078 µg/ml – 20 µg/ml respectively. Serum samples were diluted until the LBP or Haptoglobin concentrations were in the range of these kits.

### Human faecal bacterial DNA extraction and 16S rRNA amplicon sequencing

Total genomic DNA were isolated from human faecal samples using DNeasy Blood and Tissue kit (Qiagen, UK) following the manufacturer’s instructions with some modifications following the repeated bead-beating method (Yu and Morrison, 2004). The V3-V4 hypervariable region of the16S rRNA gene was amplified to generate a fragment of 460 bp using the forward primer 5′TCGTCGGCAGCGTCAGATGTGTATAAGAGACAGCCTACGGGNGGCWGCAG and reverse primer 5′GTCTCGTGGGCTCGGAGATGTGTATAAGAGACAGGACTACHVGGGTATCTAATCC (Klindworth *et al*., 2013). The Illumina overhang adapter sequences were added to the 16S rRNA gene specific primer sequences. Each 30 µl PCR reaction contained 10 ng/µl microbial genomic DNA, 0.2 µM of each primer, 15 µl of 2x Phusion Taq High-Fidelity Mix and 10.6 µl of nuclease free water. The PCR conditions were: initial denaturation 98°C for 30s; 25 cycles of 10s at 98°C, 15s at 55°C and 20s at 72°C; and 72°C for 5 minutes for final elongation. The SPRI select reagent kit (Beckman Coulter, UK) was used to purify the amplicons. The Qubit^®^ dsDNA HS Assay Kit (Life Technologies) was followed for quantification and pooling. Library preparation was carried out by Teagasc, Fermoy, Ireland on the Illumina MiSeq platform using paired-end Illumina sequencing run (2 × 250 bp).

### *APOE* Targeted Replacement mice

All experimental procedures and protocols used in this study were reviewed and approved by the Animal Welfare and Ethical Review Body (AWERB) and were conducted within the provisions of the Animals (Scientific Procedures) Act 1986 (ASPA).

Male human *APOE3* (B6.129P2-Apoe^tm2(APOE*3)Mae^ N8) and *APOE4* (B6.129P2-Apoe^tm2(APOE*4)Mae^ N8) Targeted Replacement (TR) mice homozygous for the human *APOE3* or *APOE4* gene (Taconic, Germantown, NY, US) were used in these experiments. The model was created by Dr Maeda by targeting the murine *APOE* gene for replacement with the human *APOE3* and *APOE4* allele in E14TG2a ES cells and injecting the targeted cells into blastocysts. Resultant chimeras were backcrossed to C57BL/6 for eight generations (N8). Mice were maintained in controlled environment (21°C; 12-h light–dark cycle; light from 07:00 hours) and fed a standard chow diet (RM3-P, Special Diet Services, Essex, UK) for the duration of the experiments.

### Mice genomic DNA extraction and 16S rRNA amplicon sequencing

Bacterial genomic DNA was extracted from faecal samples using a FastDNA SPIN Kit for Soil (MP Biomedicals) with three bead-beating periods of 1 min (Maukonen *et al*., 2006). Bacterial DNA concentration was normalised to 1 ng/μL by dilution with DNA elution solution (MP Biomedicals, UK) to produce a final volume of 20 μL. Normalised DNA samples were sent to the Centre of Genomic Research (Liverpool, UK) for PCR amplification of the 16S ribosomal RNA (rRNA) gene and paired-end Illumina sequencing (2 × 250 bp) on the MiSeq platform. The V4 region of the 16S rRNA gene was amplified to generate a 254 bp insert product as described previously (Caporaso *et al*., 2011). The first round of PCR was performed using the forward primer 5′‐ ACACTCTTTCCCTACACGACGCTCTTCCGATCTNNNNNGTGCCAGCMGCCGCGGTAA-3′ and the reverse primer 5′‐ GTGACTGGAGTTCAGACGTGTGCTCTTCCGATCTGGACTACHVGGGTWTCTAAT-3′, which include recognition sequences that enable a second nested PCR, using the N501f and N701r primers, to incorporate Illumina adapter sequences and barcode sequences. The use of these primers enables efficient community clustering for the length of reads obtained through Illumina sequencing, and this method also allows for high-throughput sequencing. Sequencing data were supplied in FASTQ format with adaptors already trimmed.

### Metabolomic analyses

Faecal contents were extracted from caeca and prepared for NMR analysis by mixing thoroughly 20 mg of frozen faecal material with 1 mL of saline phosphate buffer (1.9 mM Na^2^ HPO_4_, 8.1 mM NaH_2_PO_4_, 150 mM NaCl, and 1 mM TSP (sodium 3-(trimethylsilyl)-propionate-d4)) in D_2_O (deuterium oxide), followed by centrifugation (18,000g, 1 min). Supernatants were removed, filtered through 0.2 μm membrane filters, and stored at −20 °C until required.

High resolution ^1^H NMR spectra were recorded on a 600 MHz Bruker Avance spectrometer fitted with a 5 mm TCI cryoprobe and a 60 slot autosampler (Bruker, Rheinstetten, Germany). Sample temperature was controlled at 300 K. Each spectrum consisted of 128 scans of 32 768 complex data points with a spectral width of 14 ppm (acquisition time 1.95 s). The *noesypr1d* pre-saturation sequence was used to suppress the residual water signal with low power selective irradiation at the water frequency during the recycle delay (D1 = 2 s) and mixing time (D8 = 0.15 s). A 90° pulse length of 8.8 μs was set for all samples. Spectra were transformed with a 0.3 Hz line broadening and zero filling, manually phased, baseline corrected, and referenced by setting the TSP methyl signal to 0 ppm. Metabolites were identified using information found in the literature or on the web (Human Metabolome Database, http://www.hmdb.ca/) and by use of the 2D-NMR methods, COSY, HSQC, and HMBC (Le Gall *et al*., 2011) and quantified using the software Chenomx^®^ NMR Suite 7.0^TM^.

### Analysis of 16S amplicon sequencing data from humans and mice

Bioinformatics analysis of 16S amplicon sequencing data from humans and mice were performed using the Quantitative Insights Into Microbial Ecology (QIIME) v1.9.1 (Caporaso *et al*., 2010) and usearch v8.1 (Edgar, 2010) software and the following procedure. First, the paired-end reads were merged using FLASH v1.2.8 (Magoc and Salzberg, 2011), then adaptors were removed from reads using cutadapt v1.8.3 (Martin, 2011). The sequences were demultiplexed and filtered using QIIME with the split_libraries_fastq.py script, all reads with a quality score below 19 were removed. Reverse primers were removed using QIIME with truncate_reverse_primer.py. An operational taxonomic unit (OTU) table was obtained using usearch. Unique sequences were filtered (derep_fulllength) and sorted by length (sortbylength) with a length of 373–473 nt for V3-V4 region and a length of 237–271 nt for V4 region. After singleton removal (sortbysize), the remaining sequences were clustered into OTUs at a default 97% sequence identity (cluster_otus) and filtered for chimeras against the ChimeraSlayer reference database (uchime_ref) (Haas *et al*., 2011). All sequences were mapped against this database (usearch_global) to generate an OTU table. Classification of representative sequences for each OTU was carried out using mothur v1.36.1 (Schloss *et al*., 2009) against the 16S rRNA reference of Ribosomal Database Project (RDP) database trainset 14 (Cole *et al*., 2009). To ensure an even sampling depth, we used QIIME to generate rarefied OTU tables with single_rarefaction.py script and compute alpha diversity metrics (chao1, phylogenetic diversity, Shannon’s diversity index, evenness) with alpha_rarefaction script and beta diversity (weighted Unifrac, unweighted Unifrac and Bray-curtis distances) with beta_diversity.py script.

### Statistical analysis

Statistical analysis was carried out using R v.3.5.1 software packages (R Core Team, 2016). The significant differences in clinical measures, alpha diversity and abundances of each taxonomic unit between two or more groups were detected using Mann-Whitney U test or Kruskal-Wallis H test with Dunn’s multiple comparison test, respectively. The *p*-values were corrected for multiple testing by Benjamini–Hochberg correction to control false discovery rate. Differences in beta diversity were determined using permutational multivariate analysis of variance (PERMANOVA) (R package *vegan*).

Multivariate statistical analysis (Sparse PLS Discriminant Analysis (sPLS-DA) and MSEA) of the ^1^H NMR data was carried out using the MetaboAnalystR 1.0.0. Co-inertia analysis (COIA) was used to investigate the relationships between the faecal metabolome and the composition of microbiota at OTU level using the co-inertia function (R package *ade4* (Dray and Dufour, 2007)). Only OTUs present in at least 50% of the samples were used in COIA analysis. Overall similarity in the structure between two datasets were measured by RV-coefficient. The significant of the RV-coefficient was tested using the Monte-Carlo permutation test (Moonseong and Ruben Gabriel, 1998).

## Supporting information

