## Supplementary material for "Association between Apolipoprotein E genotype and the gut microbiome composition in humans and mice"

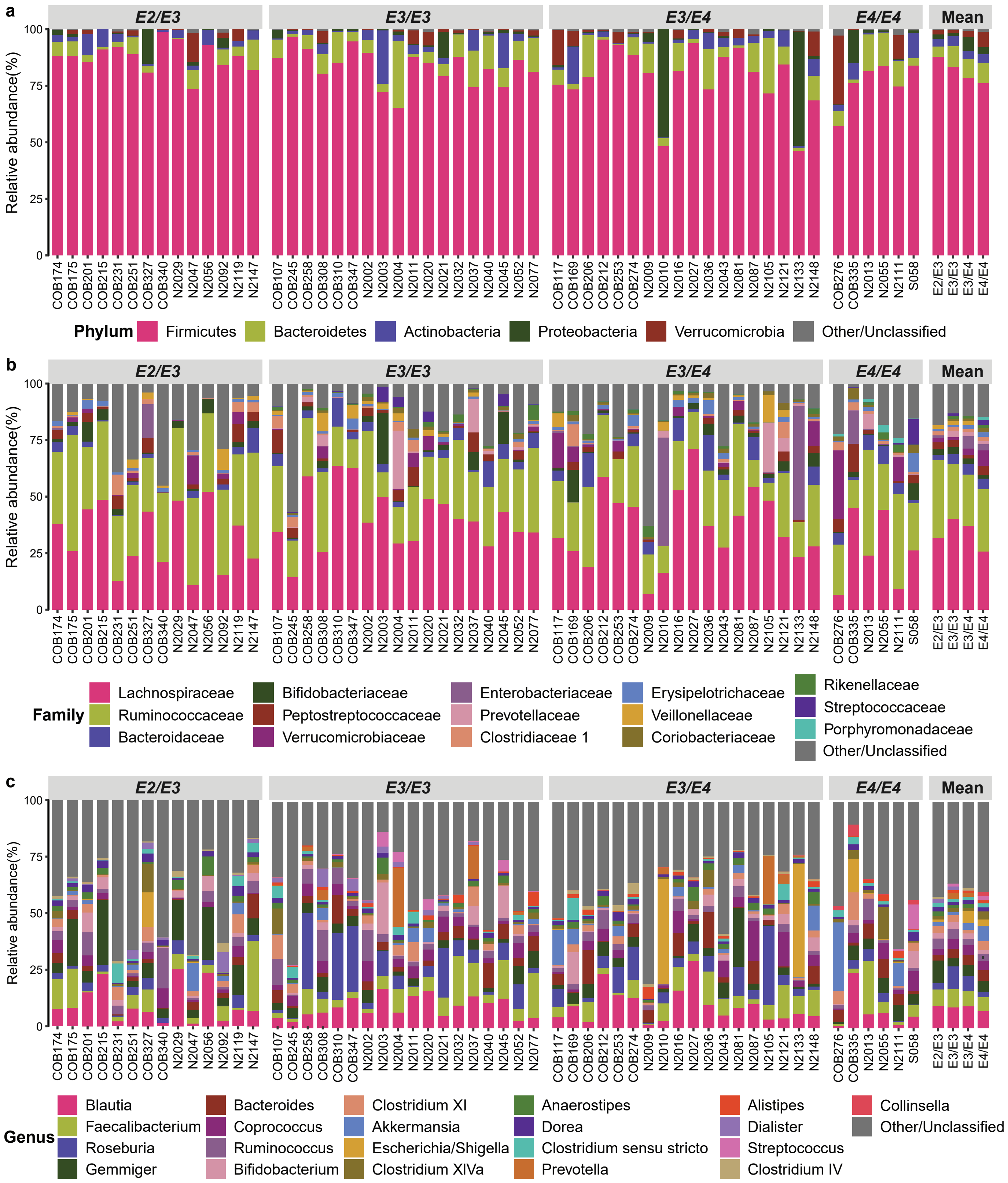


**Supplementary Figure 1.** Relative abundance of human faecal microbiota taxa at (**a**) Phylum level, (**b**) Family level and (**c**) Genus level in subjects grouped according to *APOE* genotypes.

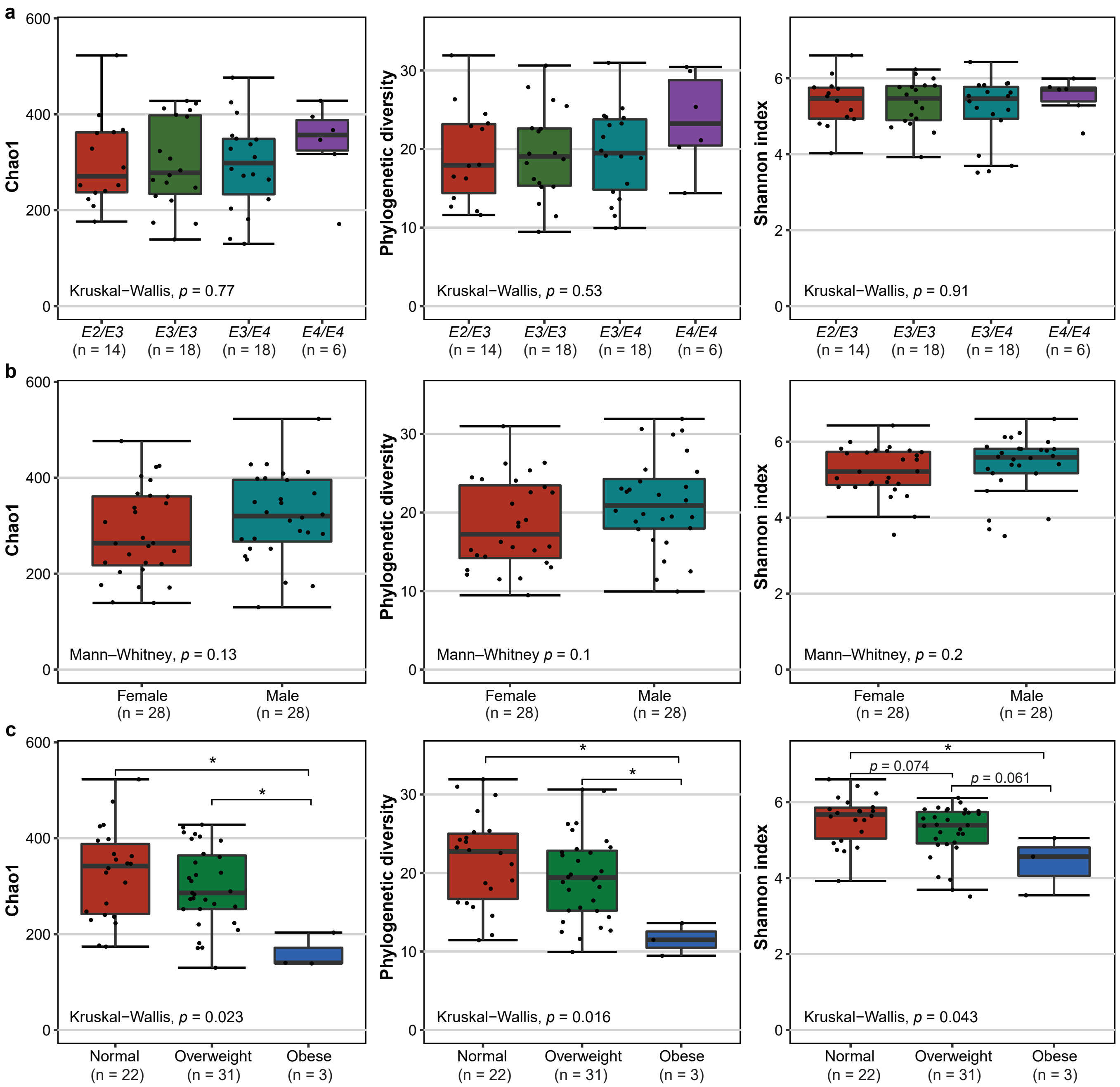


**Supplementary Figure 2.** Boxplot of bacterial alpha-diversity indices for the microbiota from human faecal samples binned according to (**a**) *APOE* genotypes (**b**) sex or (**c**) BMI categories. *P*-values were calculated by Mann–Whitney U test or Kruskal-Wallis H test followed by Dunn's multiple comparisons adjusting false discovery rate using the Benjamini-Hochberg correction. $^{*}p < 0.05$.

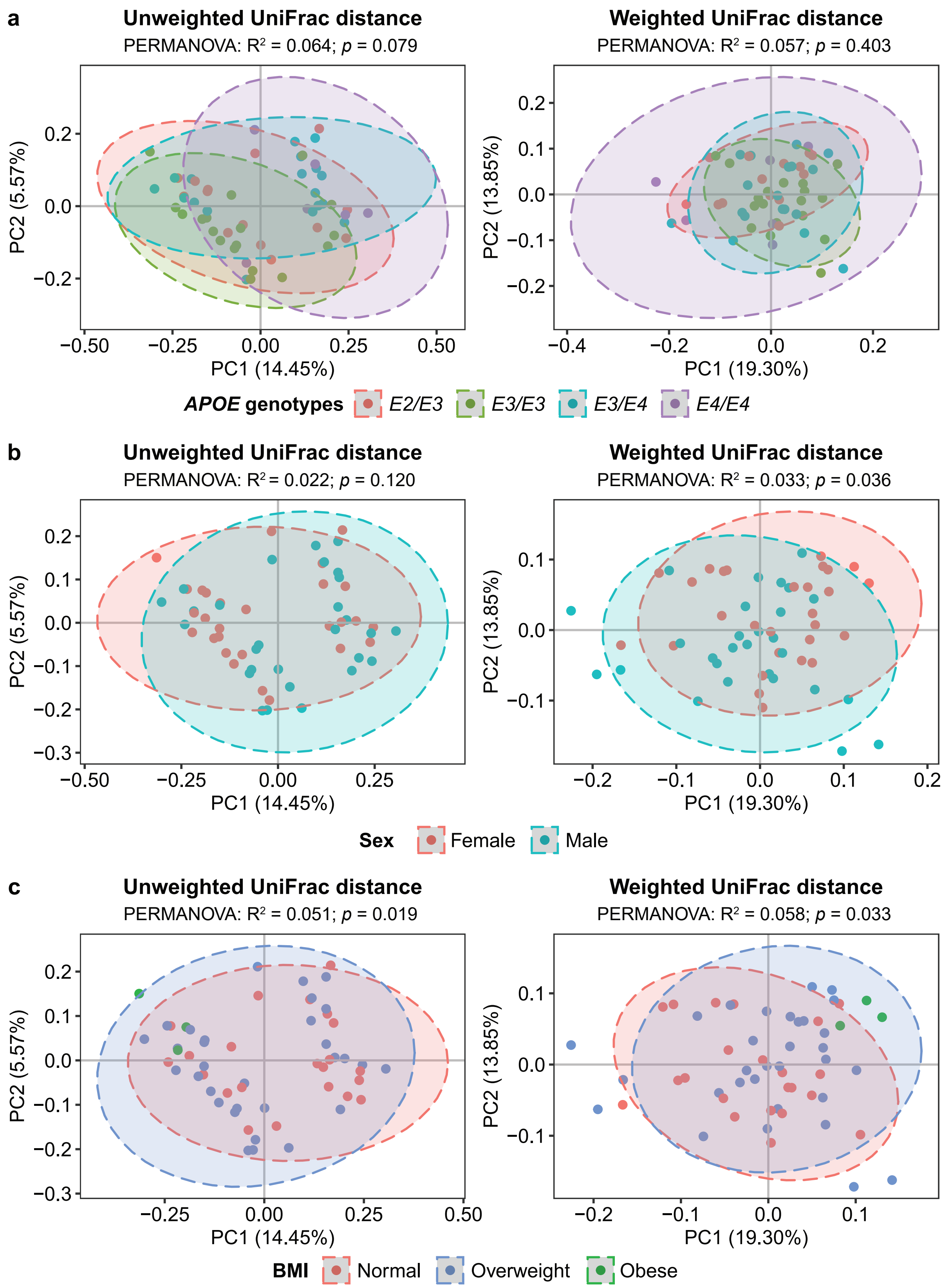


**Supplementary Figure 3.** Principle Coordinates Analysis based on unweighted and weighted UniFrac distances of bacterial 16S rRNA amplicon sequences from human faecal samples showing gut microbiota beta diversity grouped according to (**a**) *APOE* genotypes, (**b**) sex or (**c**) BMI categories. The significant differences between groups were determined by PERMANOVA tests.

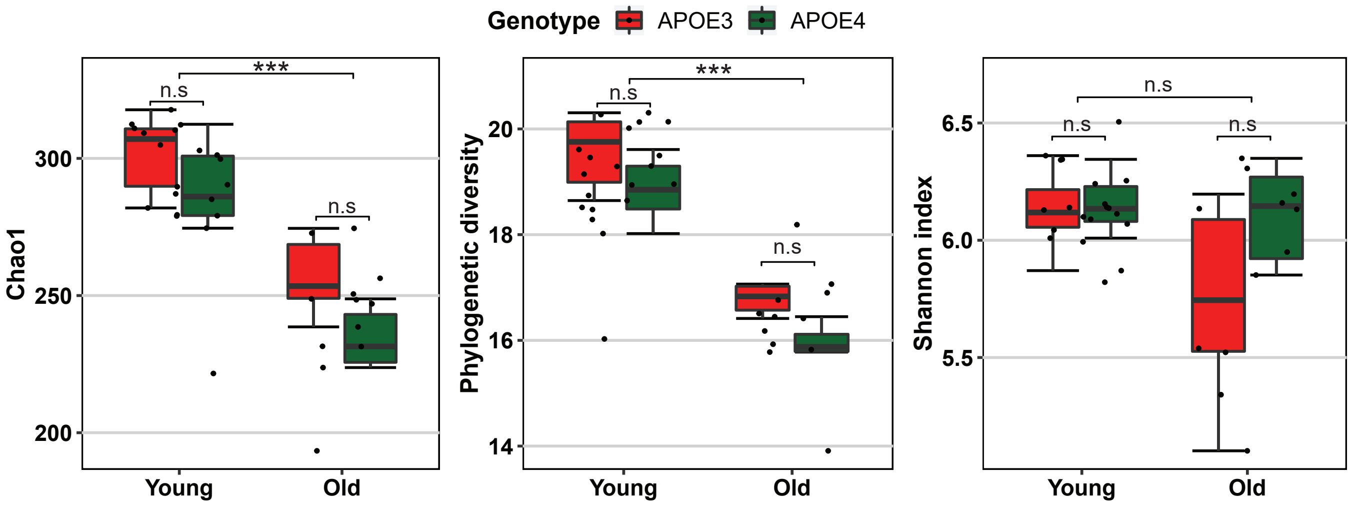


**Supplementary Figure 4.** Boxplot of bacterial alpha-diversity indices for the faecal microbiota from mice grouped according to *APOE* genotype and age. *P*-value was calculated by Mann–Whitney U test for two *APOE* genotypes or two age groups. ***$p < 0.001$; n.s, not significant.

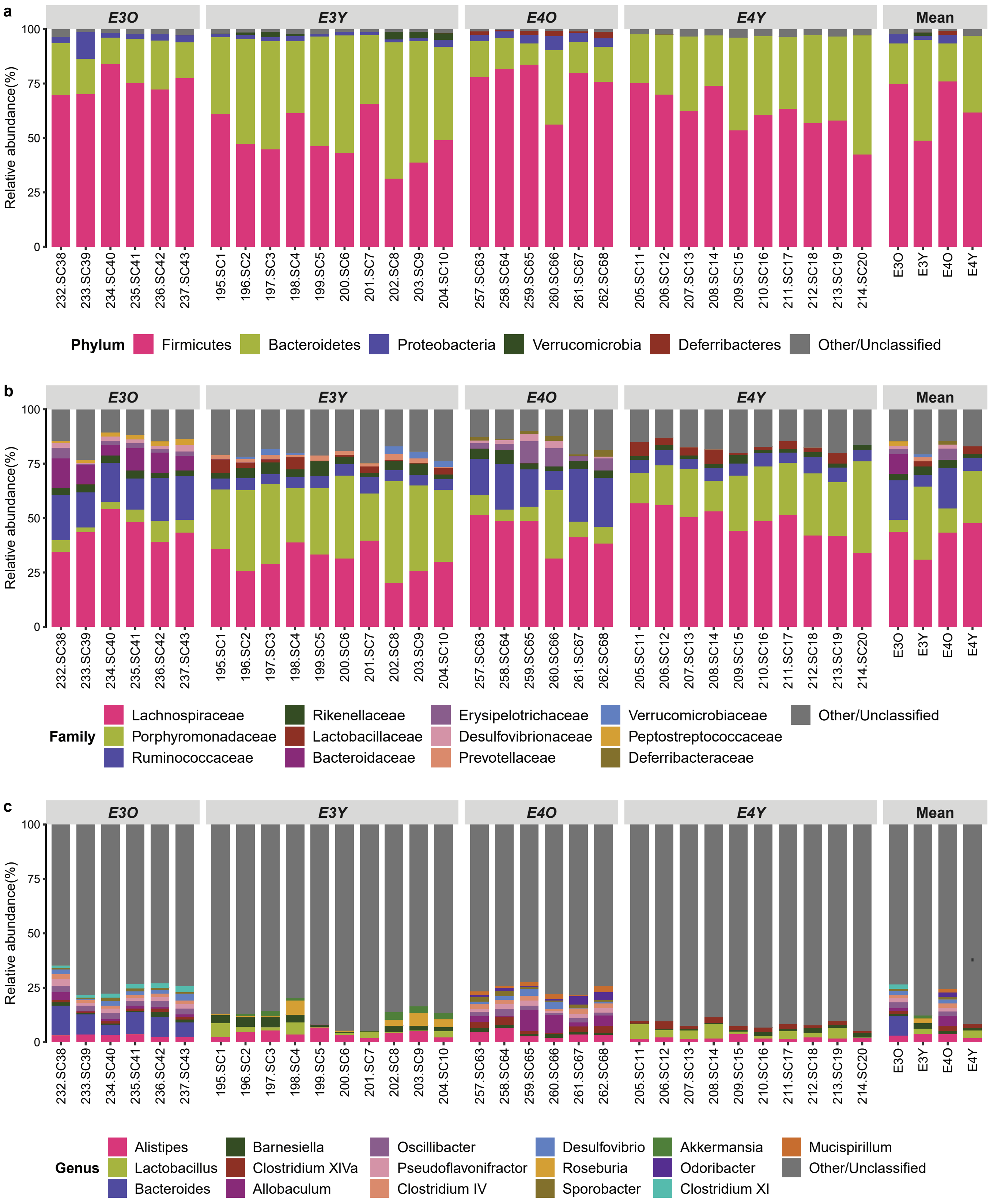


**Supplementary Figure 5.** Relative abundance of mouse faecal microbiota taxa at (**a**) Phylum level, (**b**) Family level and (**c**) Genus level in different subjects grouped according to *APOE* genotypes and age.

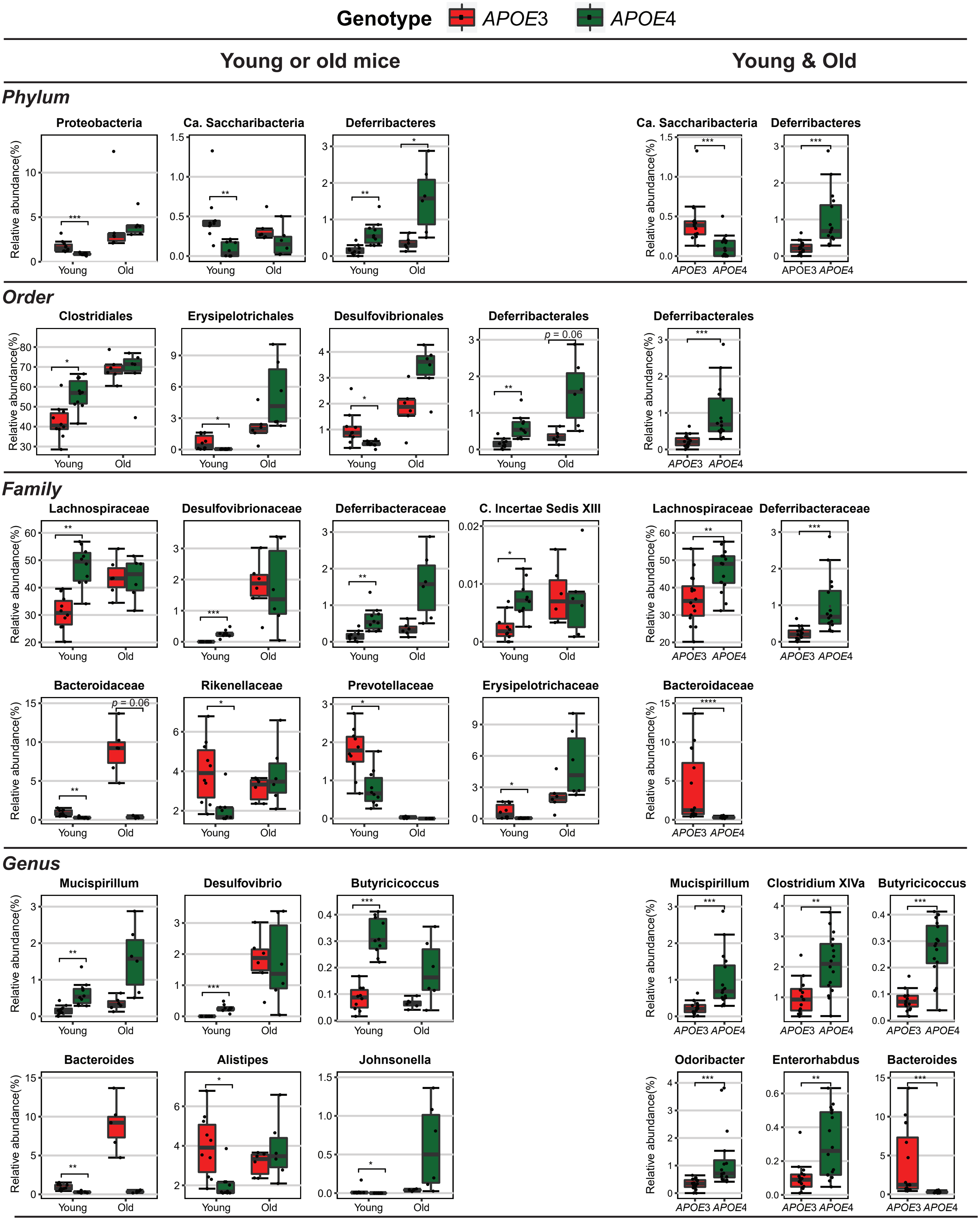


**Supplementary Figure 6.** Box plot of the relative abundance of significantly differentially abundant faecal microbiota taxa from phylum to genus associated with murine *APOE* genotypes in young mice samples, old mice samples and both age groups combined. Statistical significances between *APOE*3 and *APOE*4 were determined by the Mann–Whitney U test and were corrected for the multiple comparison using the Benjamini–Hochberg adjustment, $*p < 0.05$, $**p < 0.01$, $***p < 0.001$.

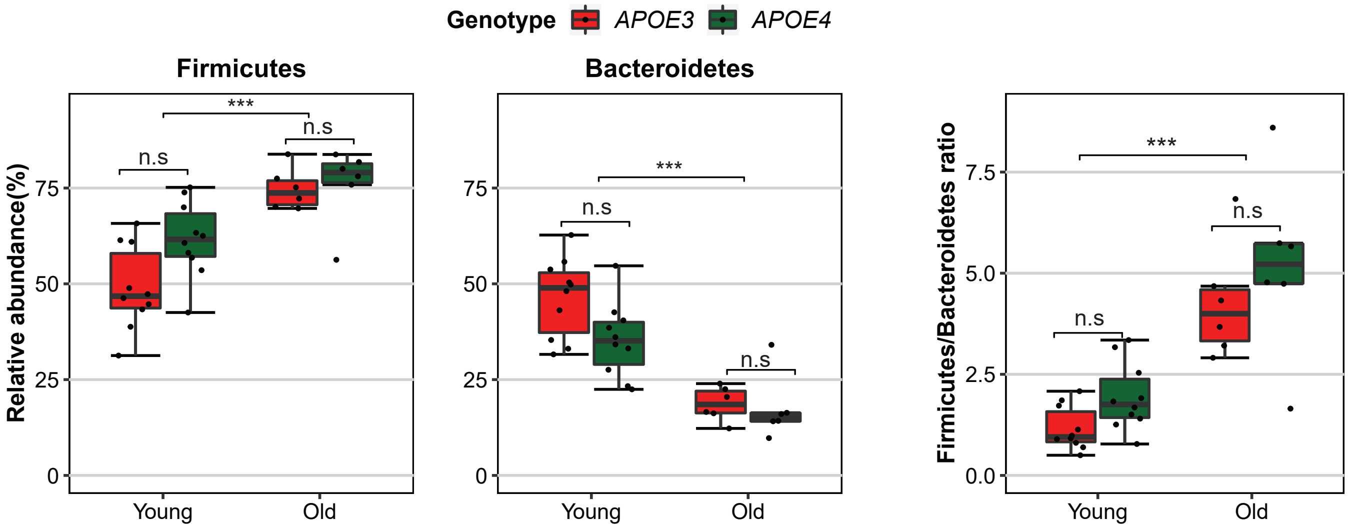


**Supplementary Figure 7.** Box plot of the relative abundance from murine faecal samples of Firmicutes, Bacteroidetes and Firmicutes/Bacteroidetes ratio according to age and *APOE* genotypes. *P*-values were calculated by Mann–Whitney U test for testing the significant difference between *APOE* genotype groups or age groups, ***$p < 0.001$; n.s, not significant.

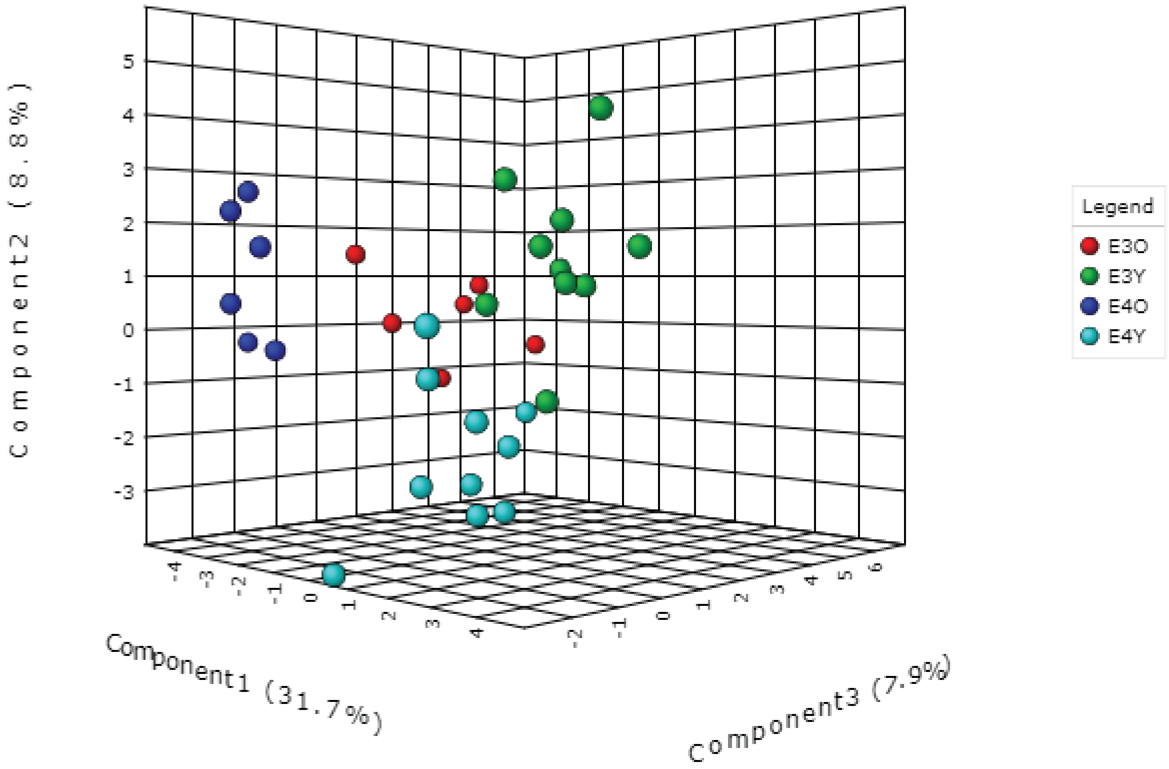


**Supplementary Figure 8.** Sparse PLS Discriminant Analysis (sPLS-DA) shows a trend for microbiota separation according to age and *APOE* genotype.

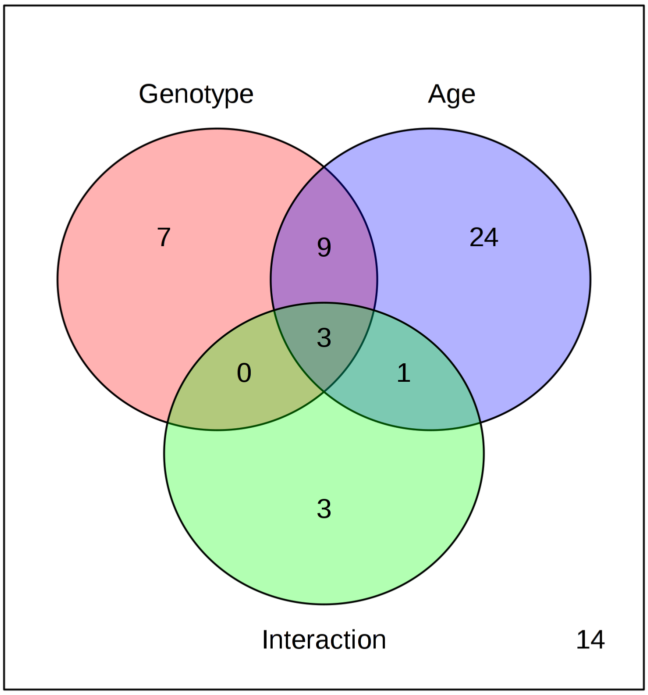


**Supplementary Figure 9.** Venn diagram summary of statistically significant metabolites according to *APOE* genotype and age were conducted by two-way ANOVA with False Discovery Rate (FDR) correction

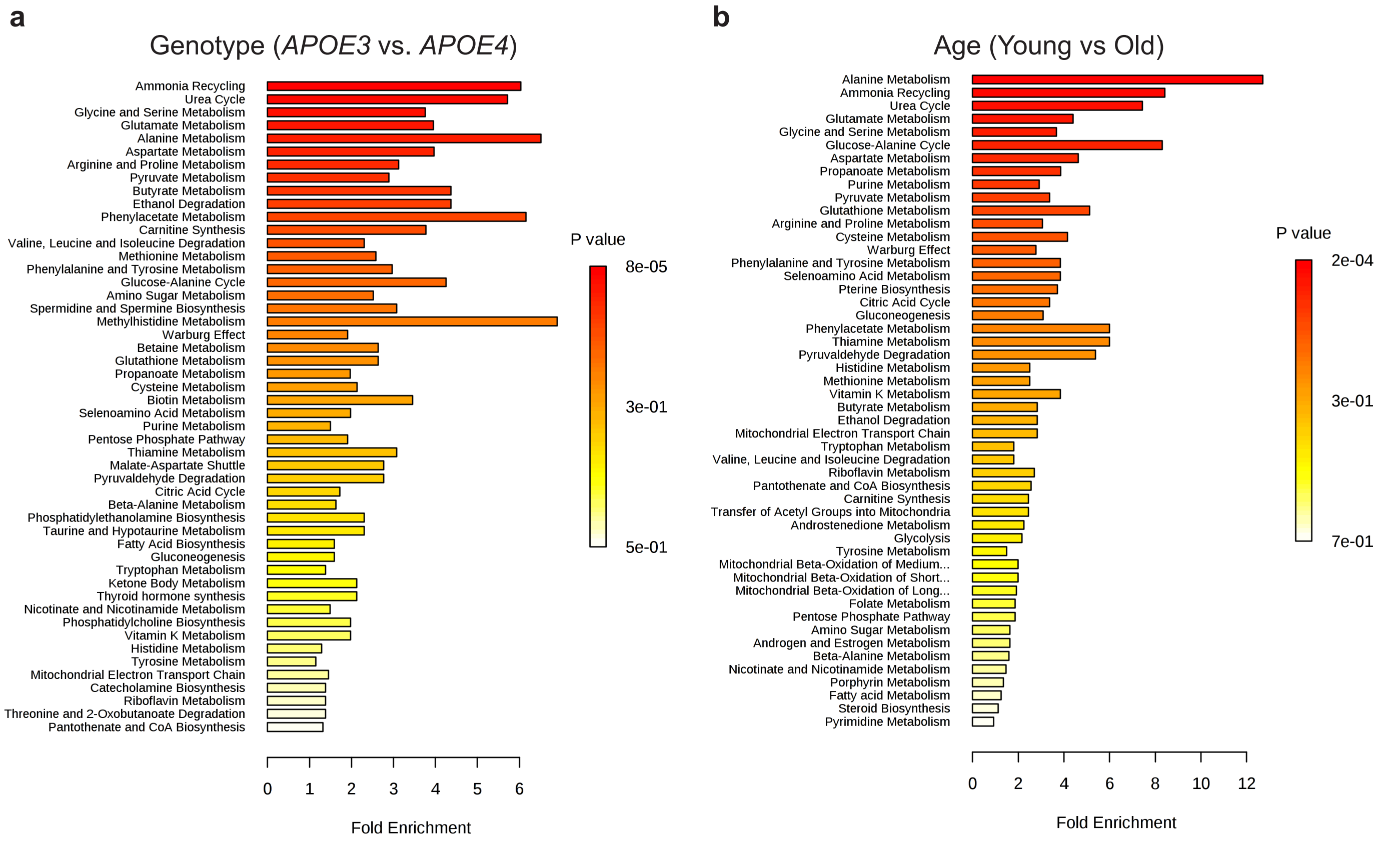


**Supplementary Figure 10.** Metabolites Set Enrichment associated to (**a**) *APOE* genotype and (**b**) age for faecal metabolites.

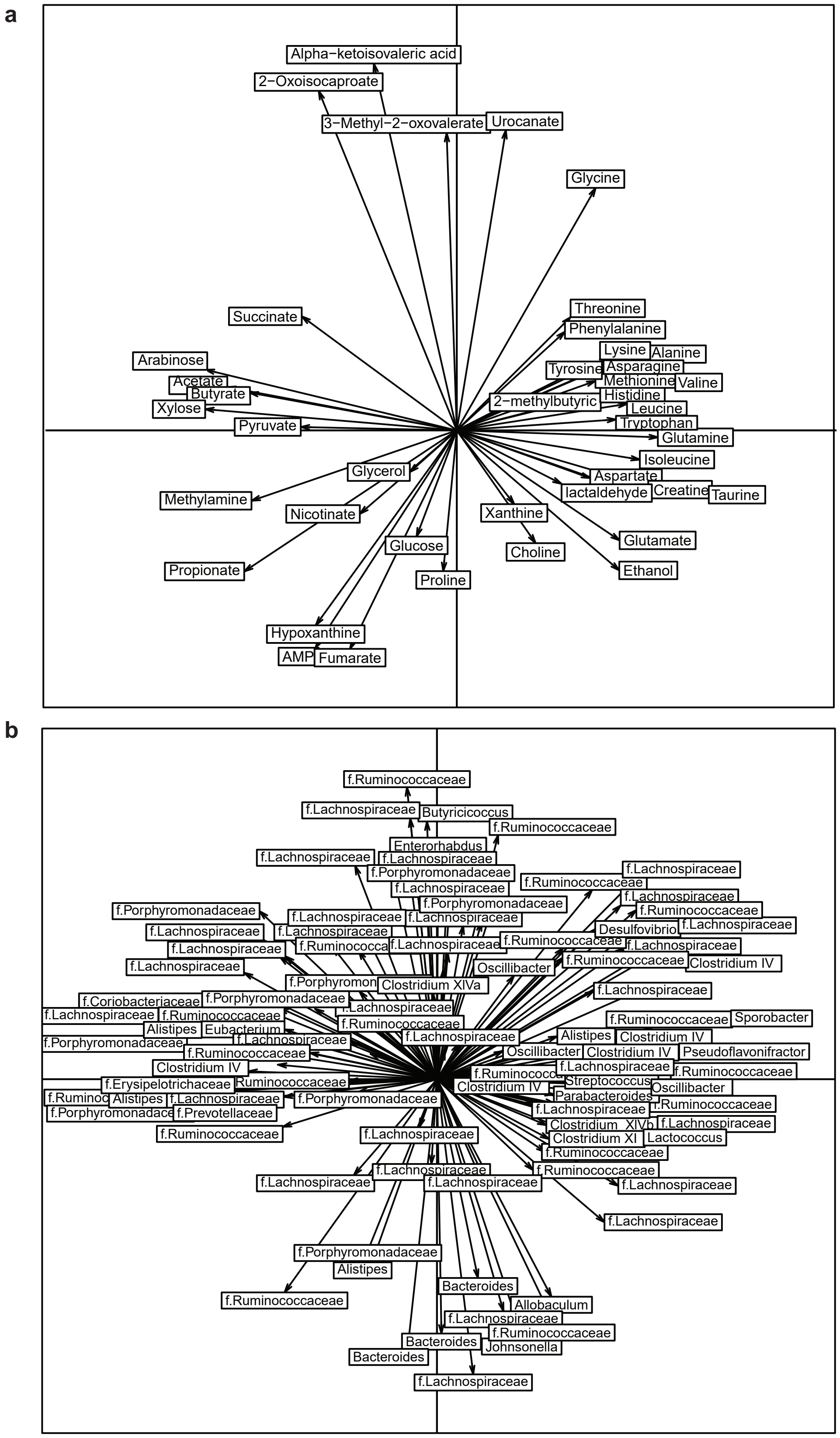


**Supplementary Figure 11. Co-inertia analysis (COIA).** Projection of (**a**) metabolites and (**b**) bacterial OTUs labelled with either genus name or family (f.) names for unclassified genera on the first two COIA axes, with the length of arrow indicating the strength of the correlation. Only metabolites and OTUs that are highly correlated with corresponding COIA axes are shown.
